# Enhanced Subjective Performance Achievement in Wind Instrument Playing through Positive Memory Recall: Effects of Sympathetic Activation and Emotional Valence

**DOI:** 10.1101/2024.12.12.628097

**Authors:** Aiko Watanabe, Sotaro Kondoh, Tomohiro Samma, Shinya Fujii

## Abstract

Controlling physiological and psychological states before a performance is essential for professional musicians to realize their full potential. However, the characteristics of the optimal pre-performance state remain unclear. While an increase in sympathetic nervous system activity is typically observed before performance, when associated with anxiety, it can degrade the performance quality. This study examined whether recalling positive autobiographical performance memories enhances subjective performance achievement, accompanied by increased emotional arousal, valence, and autonomic nervous system activity. Thirty-six professional wind instrument players participated in the study. Prior to performing musical pieces, participants engaged in one of three conditions: (1) recalling positive autobiographical memories, (2) recalling negative autobiographical memories, or (3) imagining routine pre-performance activities (no-memory condition). During the memory recall phase, heart rate was measured. After each performance, participants rated their subjective arousal, valence, and performance achievement. We calculated the heart rate variability indices, specifically SD1 (reflecting parasympathetic nervous system activity) and SD2/SD1 (reflecting sympathetic nervous system activity). The results showed that performance achievement, arousal, and valence were significantly higher in the positive than in the negative condition. Our path analysis further revealed that an increase in SD2/SD1 did not directly predict performance achievement; instead, it was associated with an increase in emotional valence, which in turn led to improved performance. These findings suggest that recalling positive performance memories activates sympathetic nervous system activity and fosters positive emotions, thereby enhancing the performance achievement of professional musicians.

## 1 Introduction

For professional musicians, controlling their psychophysiological state before a performance is crucial not only for delivering high-quality performance, but also for sustaining long-term careers. However, managing this state is often challenging because of various uncontrollable factors such as the audience, program, venue, and fellow performers (Baumeister, 1984; Burin & Osório, 2017; Furuya et al., 2021; Kenny, 2011). These factors activate the sympathetic nervous system (SNS), a division of the autonomic nervous system (ANS), leading to physiological responses, such as increased heart rate, elevated respiratory rate, and sweating. These physiological responses are correlated with psychological reactions, including heightened anxiety (Abel & Larkin, 1990; Brouwer & Hogervorst, 2014; Mulcahy et al., 1990; Yoshie et al., 2009a). Such psychophysiological changes can degrade performance skills, resulting in unsatisfactory outcomes for musicians (Yoshie et al., 2009b).

Conversely, some studies indicate that SNS activation can coincide with richer emotional expressions during performance, thereby enhancing performance quality. For example, increased sympathetic activity has been observed during emotionally expressive performance (Nakahara et al., 2011). Moreover, a music competition winner demonstrated lower anxiety and superior performance despite exhibiting similar levels of sympathetic activation compared to other contestants (Yoshie et al., 2009c). These findings suggest that, even when SNS is activated, high-quality performance can be achieved by managing this activation. However, the mechanisms through which musicians regulate sympathetic activation to enhance performance quality remain unclear.

One promising approach to enhancing performance is to cultivate positive emotions during SNS activation prior to performance. Research on musical performance has shown that heart rate significantly increases, and SNS activity, closely linked to the fight-or-flight response preparing the body for action, intensifies immediately before taking the stage (Craske & Craig, 1984; Nakahara et al., 2011; Yoshie, 2008a). Previous studies have demonstrated that positive emotional experiences can balance the activity of the sympathetic and parasympathetic nervous systems (PNS) (Kop et al., 2011; Kreibig, 2010; McCraty et al., 1995). The PNS is associated with rest-and-digest functions (Guyton & Hall, 2006; Saper, 2002) and can suppress SNS overactivation (Berntson et al., 1991). In contrast, negative emotional experiences are suggested to activate SNS activity, while having little effect on PNS activity (Kop et al., 2011; Kreibig, 2010; McCraty et al., 1995). Additionally, according to theories of emotion and ANS functioning, the interpretation of physiological responses leads to subjective emotional experiences (Craig, 2009; Damasio, 1996; Schachter & Singer, 1962). Therefore, even if SNS is activated, the body’s response can vary depending on whether the subjective experience is positive or negative.

The present study aimed to examine whether recalling positive emotions enhances emotional valence even when SNS is activated and whether this leads to improved performance achievement in professional musicians. Since previous research suggests that autobiographical memories influence ANS responses (Kop et al., 2011; Strohm et al., 2021), we hypothesized that recalling a positive performance memory would activate the SNS, mitigate reductions in PNS activity, increase emotional arousal and valence, and enhance performance achievement compared with recalling negative memories. Additionally, we explored whether pre-performance ANS activity directly predicts post-performance achievement or indirectly influences emotional arousal and valence during performance, which, in turn, affects achievement. We hypothesized that ANS activity would influence emotional arousal and valence, thereby enhancing performance.

## 2 Materials and methods

### 2.1 Participant

Thirty-six active Japanese professional classical wind instrumentalists (10 males, 26 females; mean age ± SD: 37.69 ± 8.79 years, range: 24–60 years) participated in this study. All participants graduated from music universities and specialized in the instruments they played during the experiment (see **Table 1**). Their professional experience ranged from 3 to 38 years (mean ± SD: 15.72 ± 8.75 years). To ensure extensive performance experience and the ability to provide expert evaluations, the following inclusion criteria were established: participants had graduated from a music university or received equivalent professional training and had at least three years of professional experience as wind instrumentalists. Participants were recruited through the internet and social media platforms.

**Table 1.** Participants’ instruments.

| Instrument | # Participants | Instrument | # Participants |
| --- | --- | --- | --- |
| Flute | 5 | Baritone Saxophone | 1 |
| Oboe | 2 | Trumpet | 4 |
| B<sup>b</sup> Clarinet | 6 | Horn | 4 |
| Bassoon | 1 | Trombone | 4 |
| Alto Saxophone | 6 | Euphonium | 3 |

All participants reported no history of cardiovascular or mental health issues and confirmed their physical and mental well-being. Ethical approval for this study was obtained from the Research Ethics Committee of the Keio University Shonan Fujisawa Campus (Approval Number 507). The study was conducted in accordance with the Declaration of Helsinki, and written informed consent was obtained from all the participants. The experiment was conducted in soundproofed music studios located in Tokyo and Aichi Prefecture, Japan, where temperature and humidity were controlled.

### 2.2 Procedure and data acquisition

The experimental procedure is illustrated in **Figure 1**. The participants began by sitting to measure the electrocardiogram (ECG) to calculate their baseline heart rate and then transitioned to a standing position. In each condition, the participants were asked to recall autobiographical memories, perform an assigned musical piece, and evaluate their own performance. The order of the conditions was counterbalanced across participants to minimize order effects. On a separate day, the participants listened to the recordings of other participants’ performances and provided objective evaluations.

**Figure 1.**
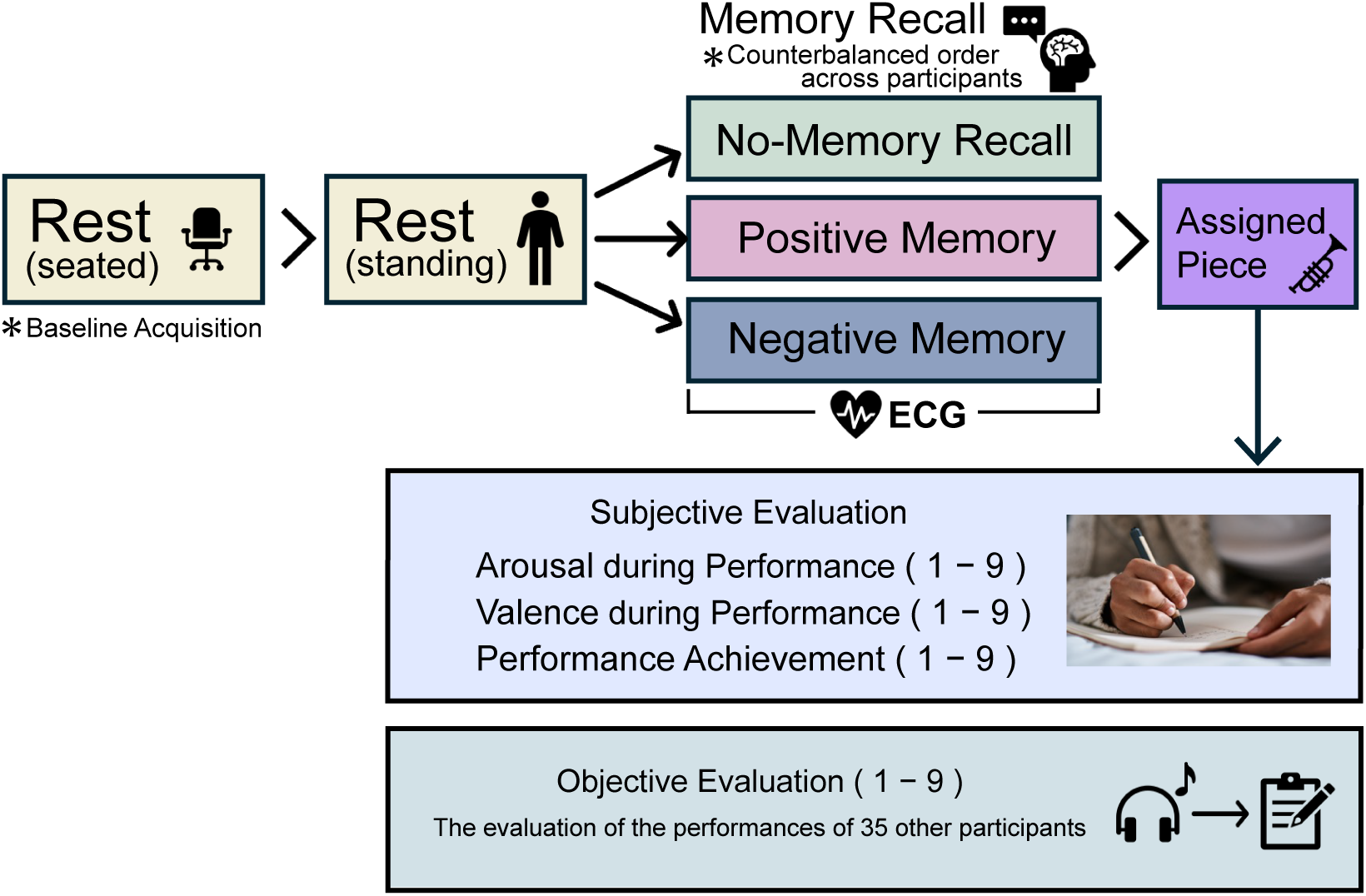
Experimental procedure. The three experimental conditions were counterbalanced across participants, and the entire procedure was repeated three times. Electrocardiogram (ECG) measurements were performed during the memory recall period. Subjective evaluations included arousal and valence, assessed retrospectively as indicators of the emotional state during the performance of the assigned piece, and performance achievement, evaluated based on the participants’ sense of accomplishment after completing the piece. On a separate day, each participant objectively evaluated the performance of other participants.

ECG data were recorded wirelessly via Bluetooth on a computer (HP ZBook Firefly, HP Inc., USA) using dedicated software (OpenSignals (r)evolution, PLUX Wireless Biosignals, Portugal) at a sampling rate of 1000 Hz. Three pre-gelled Ag/AgCl electrodes (biosignalsplux, PLUX Wireless Biosignals, Portugal) were placed on the participants’ chests to ensure accurate ECG measurements.

All performances during the session were digitally recorded on a separate computer (MacBook Retina, Apple Inc., USA) using music production software (GarageBand, Apple Inc., USA) at a sampling rate of 44.1 kHz. A USB microphone (AT2020USB+, Audio-Technica, Japan) was used to capture the performed musical pieces. The microphone was positioned 1 m from the instrument at the height of the participant’s mouth to ensure consistent and high-quality audio recording.

#### 2.2.1 Preparation and baseline measurement of ECG

Participants were instructed to abstain from alcohol the day before the experiment and avoid caffeine intake on the day of the experiment. Upon arrival at the music studio, disposable electrodes were attached to the chest to measure ECG, and ECG monitoring was initiated. A brief practice session familiarized the participants with the studio acoustics and performed with the electrodes attached, during which the sound recording levels were adjusted.

The participants sat on a chair positioned in front of a music stand and were instructed to rest quietly with their eyes open for 5 min. ECG data collected during the initial resting period served as baseline data. After an additional 5 min of standing rest, the participants began the first condition. Following each condition, the participants rested in a seated position for 5 min to allow their heart rate to return to baseline before starting the next condition.

#### 2.2.2 Memory recall and ECG measurement

Participants completed three conditions in this study: (1) positive condition: participants performed the pieces after vividly recalling their most positive performance memory; (2) negative condition: participants performed the pieces after vividly recalling their most negative performance memory; and (3) no-memory condition: participants performed the assigned piece after imagining a scenario of waiting backstage, as if preparing to perform on stage, without recalling any specific memory. The no-memory condition was designed to assess performance achievement and ANS activity without the cognitive load of memory recall, while maintaining a neutral emotional state. To facilitate effective recall during the experiment, the participants were instructed to reflect on their performance memories in advance.

ECG data recorded during the 5-minute memory recall period for each task were used for heart rate variability (HRV) analysis. Following recommendations that ECG data collection for HRV analysis should last for at least 5 min (Electrophysiology, 1996), the memory recall duration for each condition was set accordingly.

#### 2.2.3 Performed piece

In this study, an assigned musical piece was employed. Participants performed the piece using sheet music placed on a stand rather than playing from memory. The performance duration for the assigned piece was set to 5 min, and the piece was performed twice consecutively.

The assigned piece was "Ave Maria," composed by Charles Gounod in 1859. This composition combines the Latin text "Ave Maria" with a melody accompanied by J.S. Bach’s Prelude No. 1 from The Well-Tempered Clavier, Book 1 (BWV 846). We selected this piece because of its accessibility to various wind instruments, manageable range, and low technical difficulty.

Musical tempo and dynamics are known to influence cardiovascular and respiratory systems (Bernardi et al., 2006, 2009). To minimize physiological variability among participants owing to instrument-related differences, the piece’s moderate tempo and gentle dynamics were deemed appropriate. To further ensure ease of performance, participants were allowed to select the key and were instructed to prepare the piece at a performance tempo of 72 beats per minute (bpm).

#### 2.2.4 Subjective evaluation

At the end of each condition, participants provided subjective evaluations of their performance using a 9-point Likert scale to assess performance achievement (1 = did not perform to potential at all, 9 = performed to potential extremely well).

The Affect Grid (Russell et al., 1989; Takada & Yukawa, 2014) was employed to retrospectively evaluate emotional valence and arousal during performance. This tool allowed participants to rate their current emotional states on a 9×9 grid consisting of 81 cells. The horizontal axis represents the pleasure–displeasure dimension (emotional valence), where a score of 9 indicates high pleasure and 1 indicates low pleasure. The vertical axis represents the arousal dimension, where a score of 9 indicates high arousal, and 1 indicates low arousal.

#### 2.2.5 Objective evaluation

To evaluate whether the performances conveyed the intended emotions and qualities to the audience, the participants provided objective assessments of other musicians’ renditions of the assigned pieces. For this purpose, the first 20 s of each participant’s recording were extracted and edited into 60-second clips. These clips were counterbalanced across the three experimental conditions to minimize the order effects. Evaluations were conducted using Google Forms, with assessment sessions scheduled at least one month after the experiment. Participants were instructed to use their own computers and headphones in a quiet environment.

Initially, the participants were asked to set their computer volume to 25% of the maximum level and were provided with a demo sound to facilitate accurate volume adjustment. While listening to the demo, the participants adjusted the volume to a comfortable and clear level. A headphone check test was administered to confirm proper headphone use (Woods et al., 2017). After these preparations, the participants rated each of the three performances from the other 35 participants using a 9-point scale.

### 2.3 Analysis

#### 2.3.1 Preprocessing of ECG

ECG data were exported from the OpenSignals (r)evolution software and processed using the Kubios HRV Premium software, version 4.1.0 (Kubios, Finland). During preprocessing, the automatic noise detection setting was configured to "Medium," and the beat correction setting was set to "Automatic correction" to address artifacts. The R-R interval data were subsequently resampled at 4 Hz and any abnormal heart intervals were interpolated using a third-order spline, as recommended in a previous study (Tarvainen et al., 2014).

#### 2.3.2 HRV calculation

We utilized a nonlinear analysis of HRV, which is considered less affected by respiration (Toichi et al., 1997; Nakagawa, 2016; Mishima et al., 2021), given that this study involved wind instrument players who controlled their respiration while performing musical pieces. Specifically, we employed the Poincaré plot, a nonlinear analysis method. The Poincaré plot displays the *n*th R-R interval on the x-axis and the (*n+1*)th R-R interval on the y-axis, providing a visual representation of the HRV. Poincaré plots were generated from the preprocessed R-R interval data (**Figure 2**).

**Figure 2.**
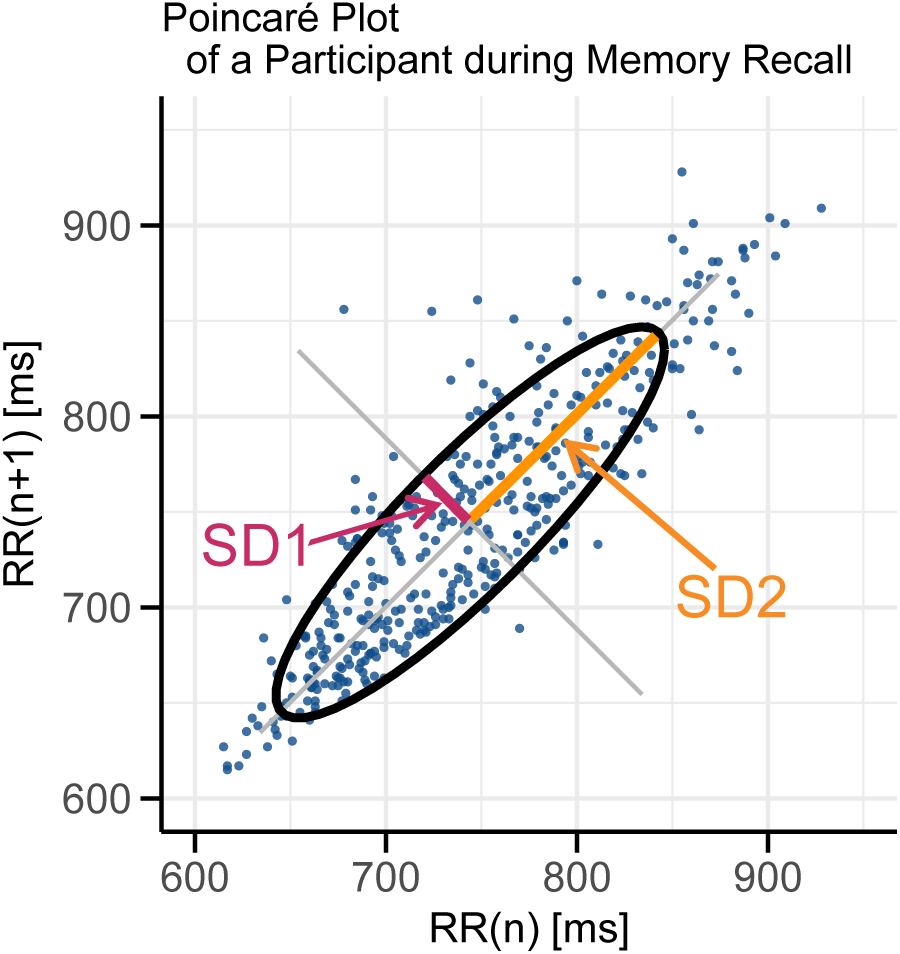
Example of Poincaré plot for a single participant. Each point represents a pair of consecutive R-R intervals: the x-axis shows the *n*th R-R interval and the y-axis shows the subsequent R-R interval. SD1 (pink line), the standard deviation along the short axis of the Poincaré plot, represents short-term fluctuations in R-R intervals and reflects parasympathetic activity. SD2 (orange line), the standard deviation along the long axis of the Poincaré plot, representing long-term variability influenced by both sympathetic and parasympathetic activities. The ellipse illustrates the overall distribution of these intervals defined by the SD1 and SD2 values.

From the Poincaré plot, we calculated SD1 and SD2, as follows:

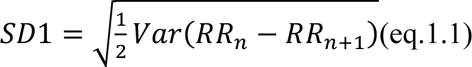

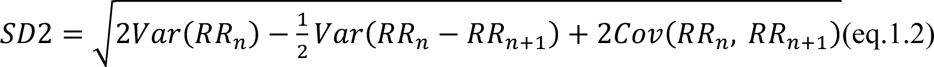

where *Var*(*RR*_*n*_) represents the variance of the *n*th R-R intervals, *Var*(*RR*_*n*_ − *RR*_*n*+1_) represents the variance of the differences between the consecutive *n*th and (*n+1*)th R-R intervals, and *Cov*(*RR*_*n*_, *RR*_*n*+1_) represents the covariance between the consecutive *n*th and (*n+1*)th R-R intervals. In this study, we analyzed HRV from a 5-minute duration of memory recall for each condition set prior to performance. We examined the SD2/SD1 ratio and SD1 as changes from the baseline to account for individual differences at rest.

SD1, the standard deviation along the short axis of the Poincaré plot, represents short-term fluctuations in R-R intervals and reflects parasympathetic activity. Short-term variability such as respiratory sinus arrhythmia is typically observed in the relaxed state. In contrast, SD2, the standard deviation along the long axis of the Poincaré plot, represents long-term fluctuations in R-R intervals and is influenced by both sympathetic and parasympathetic activities. Long-term variability increases when sympathetic activity is dominant. The SD2/SD1 ratio indicates the proportion of long-to short-term variability in R-R intervals and serves as an indicator of sympathetic activity (Brennan et al., 2001; Guzik et al., 2007; Lerma et al., 2003). According to previous studies, SD1 correlates with high-frequency (HF) components of HRV, which reflects parasympathetic activity, whereas the SD2/SD1 ratio correlates with the low-frequency to high-frequency (LF/HF) ratio, an indicator of sympathetic dominance (Goit et al., 2016; Hsu et al., 2012).

#### 2.3.3 Statistics

To evaluate the effects of memory recall on performance, we analyzed whether subjective performance achievement differed among conditions using the Friedman test, followed by Dunn’s test for multiple comparisons between the conditions. The same statistical tests were applied to emotional arousal and valence to determine whether these indicators varied across the conditions. To examine whether SD2/SD1 and SD1 varied across conditions, we constructed two linear mixed-effects models: one with SD2/SD1 as the response variable, and the other with SD1. In both models, the condition was included as a fixed effect, and participant ID as a random effect. The linear mixed-effects model was run using the ‘lme4’ and ‘lmerTest’ packages in R (Bates et al., 2015; Kuznetsova et al., 2017). The model formulas are as follows:

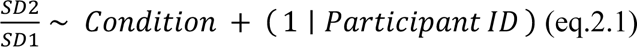

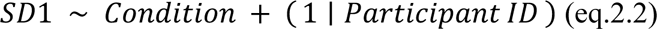

Using the positive condition as a reference, we quantitatively evaluated the deviations in the negative and no-memory conditions. Additionally, the negative condition was used as a reference to assess differences between the positive and no-memory conditions. Model fit was assessed by calculating the marginal R^2^ (R_m_^2^) and conditional R^2^ (R_c_^2^) using the ‘partR2’ package in R (Stoffel et al., 2021). R_m_^2^ represents the explanatory power of the fixed effects alone, whereas R_c_^2^ reflects the explanatory power of both the fixed and random effects.

To explore the relationships between SD2/SD1, SD1, arousal, valence, and performance achievement, we conducted a path analysis using structural equation modeling (SEM). In the model, performance achievement was predicted by valence and arousal; valence and arousal were in turn predicted by SD2/SD1 and SD1. Additionally, performance achievement was directly predicted by SD2/SD1 and SD1. Data from all conditions were combined for this analysis without accounting for repeated measurements. SEM was performed using the ‘lavaan’ package in R (Rosseel, 2012).

Additionally, the Friedman test was used to analyze whether objective evaluations by other participants differed among the three conditions. Finally, Spearman’s rank correlation coefficient was calculated to investigate the relationship between the subjective evaluations of performance in the three conditions and objective evaluations by other participants.

For all statistical tests, the significance level (α) was set at 0.05. P-values for multiple comparisons were adjusted using the Bonferroni method and are denoted as *p_b_* in the Results section.

## 3 Results

### 3.1 Performance achievement, arousal, and valence

The mean performance achievement scores were 6.06 ± 1.59 (mean ± SD) for the positive condition, 5.03 ± 1.48 for the negative condition, and 5.81 ± 1.51 for the no-memory condition (**Figure 3A**). The Friedman test revealed a significant difference in the performance achievement scores among the conditions (*χ²(2)* = 9.09, *p* = 0.011). Post-hoc comparisons using Dunn’s test indicated that the positive condition had a significantly higher achievement score than the negative condition (*z* = 3.10, *p_b_* = 0.003), and the no-memory condition also scored higher than the negative condition (*z* = 2.34, *p_b_* = 0.029). However, there was no significant difference between positive and no-memory conditions (*z* = 0.76, *p_b_* = 0.670).

**Figure 3.**
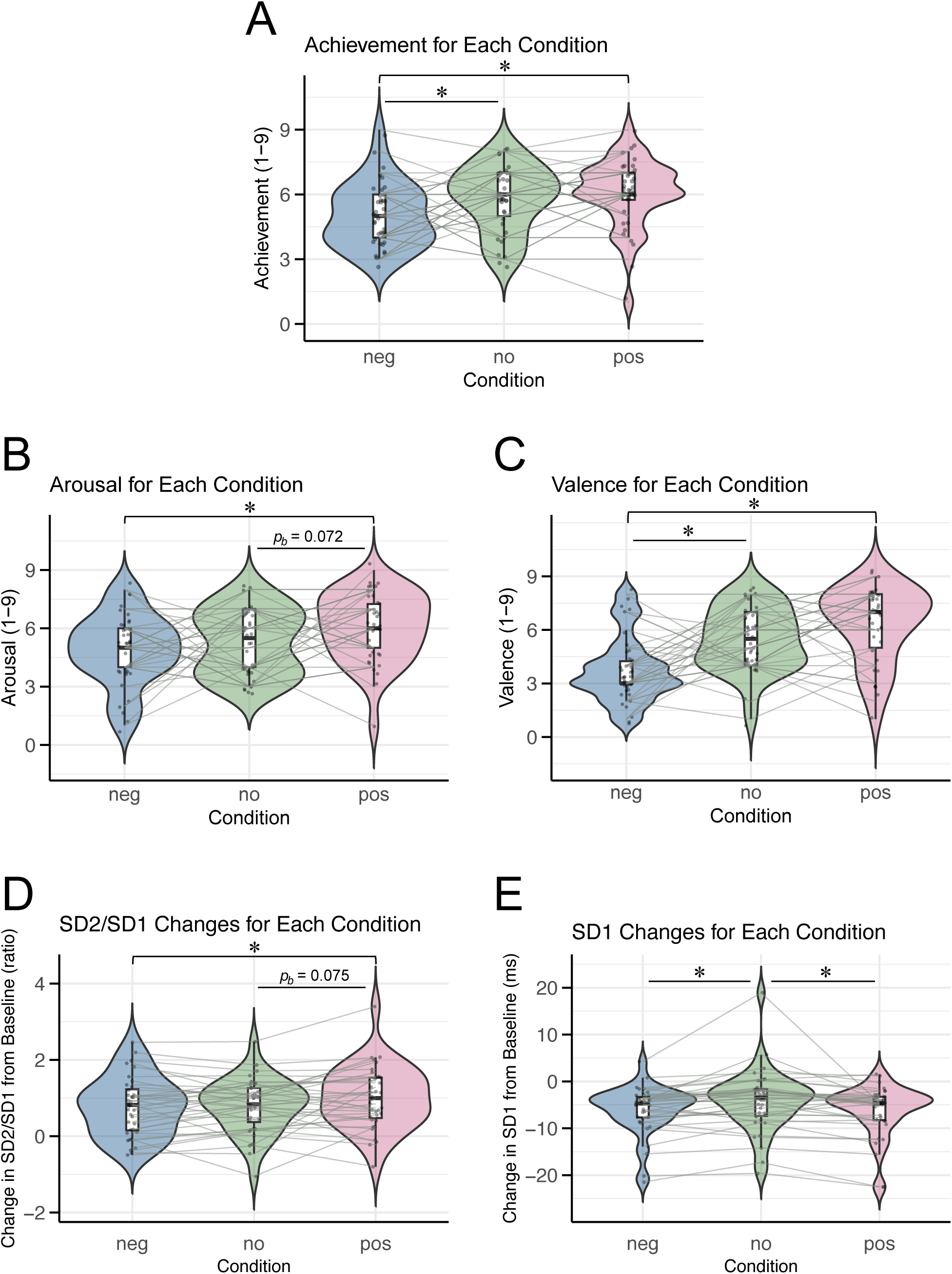
Violin plots illustrating subjective ratings and ANS indices for each condition. Each dot represents an individual participant, and asterisks (“*”) indicate significant differences between conditions (\**p_b_* < 0.05). Condition labels: “neg” = negative condition, “no” = no-memory condition, “pos” = positive condition. **(A)** Performance achievement. **(B)** Arousal. **(C)** Valence. **(D)** Changes in SD2/SD1 from the baseline. **(E)** Changes in SD1 from baseline.

The mean arousal scores were 6.06 ± 1.76 for the positive condition, 4.89 ± 1.83 for the negative condition, and 5.28 ± 1.68 for the no-memory condition (**Figure 3B**). The Friedman test showed a significant difference in arousal scores across conditions (*χ²(2)* = 7.89, *p* = 0.019). Dunn’s post hoc analysis revealed a significant difference between the negative and positive conditions (*p_b_* = 0.012). However, no significant differences were found between the negative and no-memory conditions (*p_b_* = 0.739) or between the no-memory and positive conditions (*p_b_* = 0.072).

The mean valence scores were 6.31 ± 2.10 in the positive condition, 3.78 ± 1.87 in the negative condition, and 5.47 ± 1.76 in the no-memory condition (**Figure 3C**). The Friedman test indicated highly significant differences in valence scores across the conditions (*χ²(2)* = 18.38, *p* < 0.001). Post-hoc analysis using Dunn’s test identified significant differences between the negative and positive conditions (*p_b_* < 0.001) and between the negative and no-memory conditions (*p_b_* = 0.001). By contrast, no significant difference was observed between the no-memory and positive conditions (*p_b_ =* 0.174).

### 3.2 SD2/SD1 and SD1

The mean changes in the SD2/SD1 ratio (reflecting SNS activity) from baseline were 1.00 ± 0.82 (mean ± SD) in the positive condition, 0.77 ± 0.77 in the negative condition, and 0.80 ± 0.71 in the no-memory condition (**Figure 3D**). Linear mixed-effects modeling revealed that SD2/SD1 was significantly higher in the positive condition than in the negative condition, whereas no significant difference was observed between the positive and no-memory conditions (**Table 2**). The model fit was assessed with R²m = 0.02 and R²c = 0.75.

**Table 2.** Pairwise comparisons for the conditions in SD2/SD1 using the linear-mixed regression models.

| Compared Conditions | $\beta$ | *SE* | *df* | *t-value* | *p<sub>b</sub>* |
| --- | --- | --- | --- | --- | --- |
| Negative – Positive | -0.24 | 0.09 | 70 | -2.57 | 0.037 |
| No-memory – Positive | -0.21 | 0.09 | 70 | -2.29 | 0.075 |
| No-memory – Negative | 0.03 | 0.09 | 70 | 0.29 | 1.00 |

The mean changes in SD1 (reflecting PNS activity) from baseline were −6.25 ± 5.52 ms in the positive condition, −6.09 ± 5.20 ms in the negative condition, and −4.16 ± 6.64 ms in the no-memory condition (**Figure 3E**). Linear mixed-effects modeling revealed that SD1 was significantly higher in the no-memory condition than in both the positive and negative conditions, whereas no significant difference was observed between the positive and negative conditions (**Table 3**). The model fit was assessed with R²m = 0.03 and R²c = 0.83.

**Table 3.**
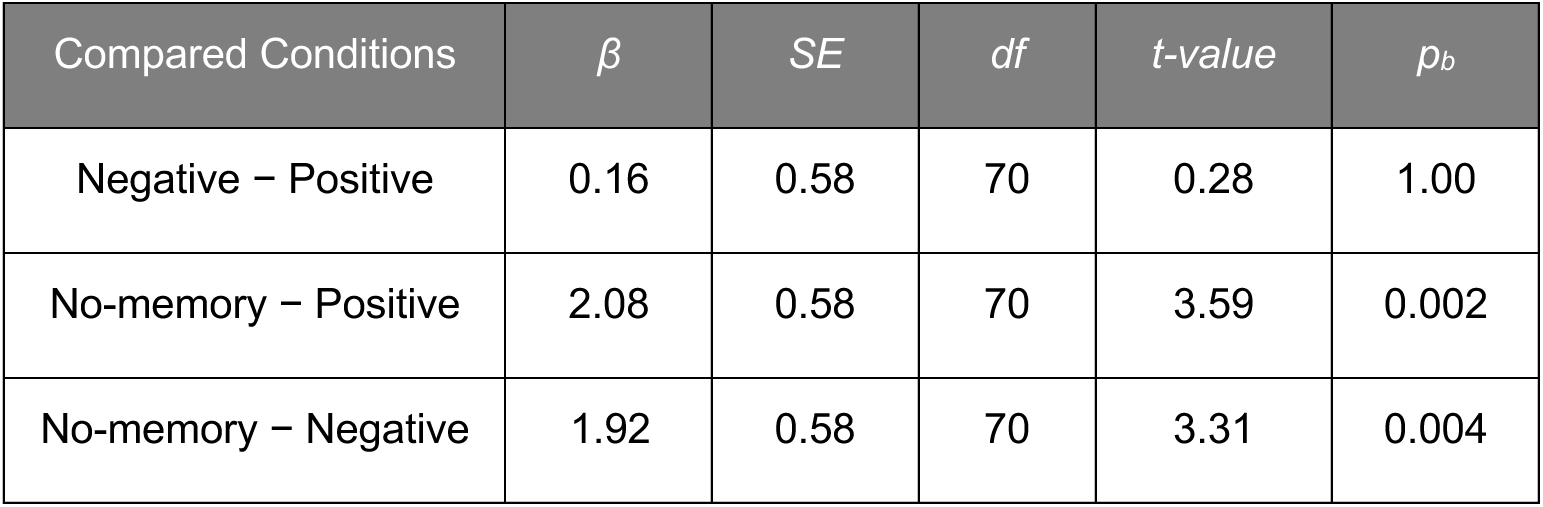
Pairwise comparisons for the conditions in SD1 using the linear-mixed regression models.

### 3.3 Relationship between psychological and physiological indicators

We examined the relationships between psychological and physiological indicators using structural equation modeling (SEM). Data from all the conditions were combined without repeated measurements (**Figure 4A**). The model tested the hypothesis that performance achievement is predicted by emotional valence and arousal, valence and arousal are predicted by SD2/SD1 and SD1, and post-performance achievement is predicted by SD2/SD1 and SD1. Model fit indices indicated a good fit (Comparative Fit Index = 1.00, Tucker-Lewis Index = 1.14, Root Mean Square Error of Approximation = 0.00, and Standardized Root Mean Square Residual = 0.01).

**Figure 4.**
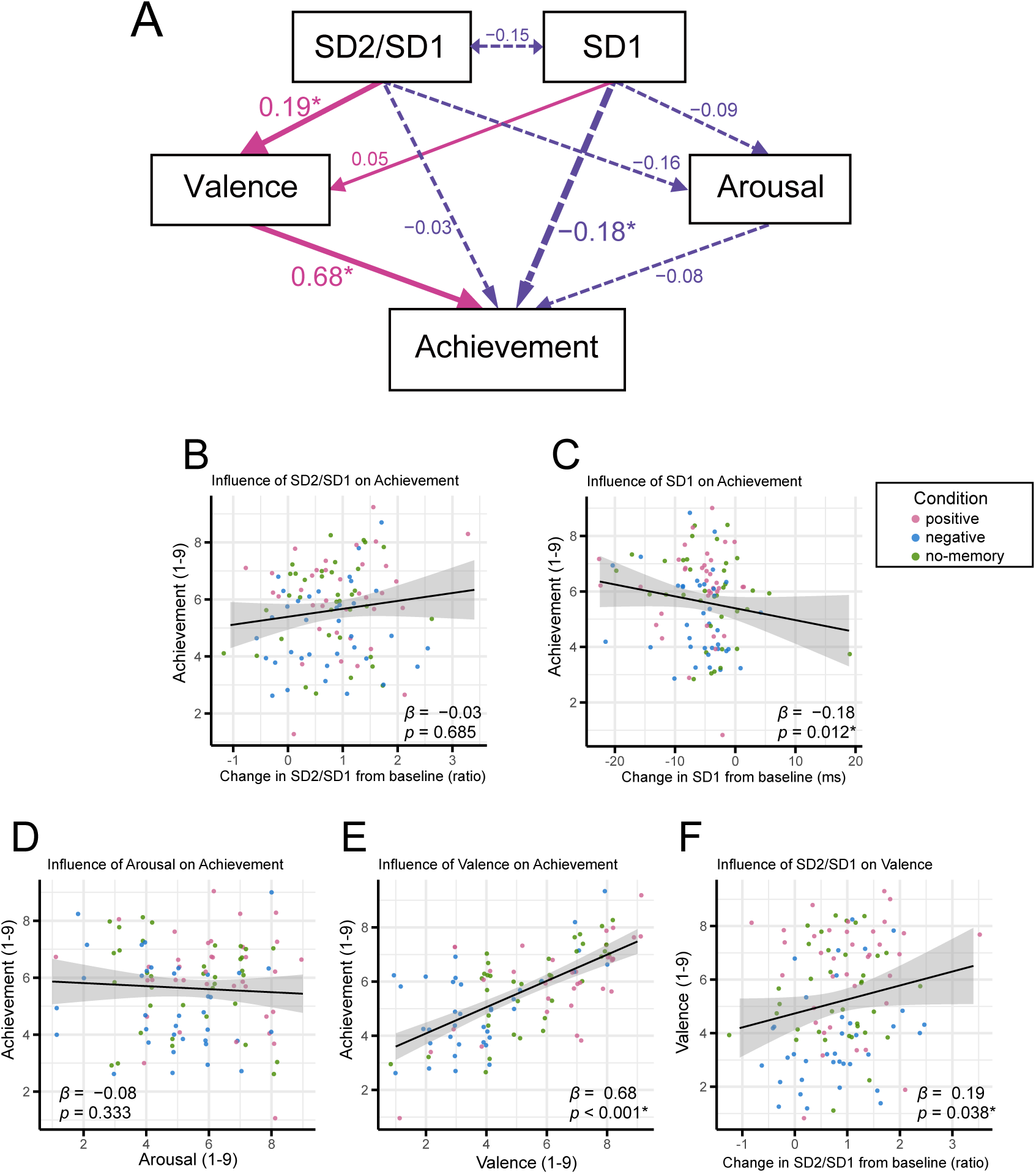
Relationship between psychological and physiological indicators. **(A)** Results of the path analysis showing how emotional valence, arousal, SD2/SD1, and SD1 influence performance achievement. Single-headed arrows represent direct predictions, whereas double-headed arrows represent correlations. Pink arrows indicate positive effects and purple (dotted) arrows indicate negative effects. The numbers next to each arrow are standardized estimates indicating the strength and direction of these effects (\**p* < 0.05). **(B-F)** Scatterplots illustrating pairwise relationships among the variables. Each dot represents an individual participant, with pink, blue, and green dots corresponding to the positive, negative, and no-memory conditions, respectively. The *β* coefficients and *p*-values shown here are consistent with those reported in the path analysis.

The scatter plots for each pairwise relationship are shown in **Figures 4B–F**. The analysis revealed that SD2/SD1 exerted a significant positive effect on valence (*β* = 0.19, *p* = 0.038; **Figure 4F**), and valence, in turn, had a significant positive effect on performance achievement (*β* = 0.68*, p* < 0.001; **Figure 4E**). Additionally, SD1 had a significant negative effect on achievement (*β* = −0.18*, p* = 0.012; **Figure 4C**).

In contrast, neither SD₂/SD₁ nor arousal significantly predicted achievement (*β* = −0.03, *p* = 0.685, *β* = −0.08, *p* = 0.333, respectively; **Figure 4B,D**). Similarly, neither SD₂/SD₁ nor SD₁ significantly predicted arousal (*β* = −0.16, *p* = 0.098 and *β* = −0.09, *p* = 0.328, respectively), and SD1 did not significantly predict valence (*β* = 0.05, *p* = 0.496). All *β* values were standardized.

### 3.4 Correlation between subjective and objective evaluations

We additionally examined whether the performance effectively conveyed the intended emotions and quality to the audience. The Friedman test revealed no significant differences between conditions (*χ²(2)* = 0.58, *p* = 0.75, **Figure 5A**). In contrast, after combining the data for all conditions, we found a significant correlation between subjective performance achievement and objective evaluations after z-scoring for both variables (Spearman’s rank correlation coefficient = 0.23, *p* = 0.016; **Figure 5B**). The 95% confidence interval, calculated using the bootstrap method, ranged from 0.06 to 0.40, confirming a weak positive correlation between subjective performance achievement and objective evaluations.

**Figure 5.**
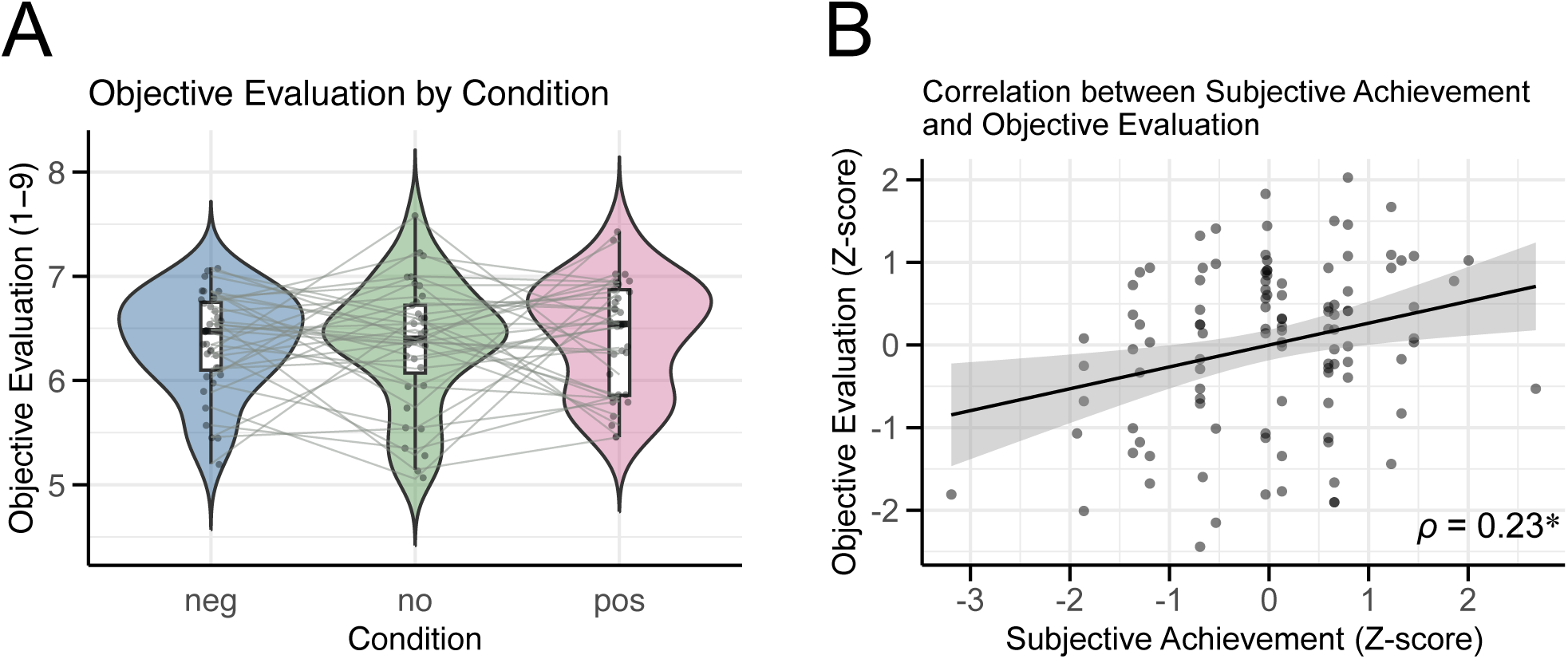
Results of the objective evaluation. **(A)** Violin plot showing the objective evaluation scores for each condition. Each dot represents the average score for each participant’s performance under these conditions. **(B)** Scatter plot illustrating the relationship between subjective achievement and objective evaluation. Each dot represents a z-score evaluation value for each participant. The regression line indicates the relationship between the two variables, and the grey band represents the standard error of the regression line. *ρ* is Spearman’s rank correlation coefficient, and a significant positive correlation was found (\**p* < 0.05).

## 4 Discussion

This study investigated whether recalling a positive performance memory could activate SNS, mitigate reductions in PNS activity, increase emotional arousal and valence, and ultimately enhance performance. We also explored whether pre-performance ANS activity directly predicted post-performance achievement or indirectly influenced emotional arousal and valence during performance, thereby affecting achievement.

### 4.1 Effects of recalling positive compared to negative performance memories

Our results revealed that subjective performance achievement was significantly higher when participants recalled positive performance memories than when they recalled negative ones (**Figure 3A**). This finding suggests that recalling positive memories can facilitate higher-level performances. In addition, emotional arousal and valence were both significantly greater in the positive condition than in the negative condition (**Figure 3B,C**), indicating that recalling positive memories not only heightens arousal, but also enhances emotional valence during performance.

We also found that the change in the SD2/SD1 ratio from baseline was significantly higher in the positive condition than in the negative condition (**Figure 3D**), suggesting a stronger SNS activation when recalling positive versus negative memories. This increase in sympathetic activity aligns with previous studies showing that recalling positive emotional experiences enhances sympathetic responses (McCraty et al., 1995), and that recalling autobiographical memories can elevate ANS responses (Strohm et al., 2021). Moreover, earlier research has established that SNS activity intensifies immediately before performance (Craske & Craig, 1984; Nakahara et al., 2011; Yoshie, 2008a), presumably preparing the body for optimal action.

Considering that the SD2/SD1 findings align with our observation of higher subjective performance achievement, arousal, and valence in the positive condition than in the negative condition, these results support the hypothesis that recalling positive performance memories, accompanied by SNS activation, enhances valence and increases arousal during performance, ultimately contributing to improved performance.

Previous studies have shown that positive emotions enhance activity in the prefrontal cortex and anterior cingulate cortex (ACC) (Ashby et al., 1999), and imagining musical performance has been observed to activate functional connectivity between the angular gyrus (AG) and ACC (Tanaka & Kirino, 2019). Integrating these findings, it is possible that recalling positive performance memories in this study facilitated processes such as emotion regulation, decision-making, and attentional control, which may have contributed to the improvement in performance achievement. Although we did not measure neural activity in this study, the observed benefits of recalling positive performance memories on performance achievement may be mediated by the underlying neural mechanisms. Future research should examine whether activation of the prefrontal cortex, ACC, and AG occurs when musicians recall positive autobiographical memories.

### 4.2 Comparisons of PNS activity between recalling positive and negative memories

We observed no significant difference in the change in SD1 between positive and negative conditions (**Figure 3E**). Given the antagonistic relationship between the sympathetic and parasympathetic nervous systems (Guyton & Hall, 2006; Saper, 2002), and considering our initial hypothesis that the positive condition would yield a higher SD2/SD1 ratio, we expected a more pronounced decrease in parasympathetic activity in the positive condition. However, the SD1 results did not support this prediction.

Previous studies have shown that positive emotions can simultaneously enhance sympathetic and parasympathetic activity (Kop et al., 2011; Kreibig, 2010; McCraty et al., 1995). This dual enhancement may explain why SD1 did not decline as much as anticipated in the positive condition despite the elevated SD2/SD1 values. In contrast, negative emotional experiences are reported to increase sympathetic activity, while exerting relatively little influence on parasympathetic activity (Kop et al., 2011; Kreibig, 2010; McCraty et al., 1995). Consequently, in the negative condition, SD1 may have decreased more proportionally in response to changes in SD2/SD1.

### 4.3 Effects of no-memory recall compared to memory-recall conditions

Interestingly, we found that performance achievement, emotional valence, and SD1 were all significantly lower in the negative memory condition than in the no-memory condition (**Figure 3A, C, E**). These results suggest that recalling negative performance memories decreases achievement, valence, and PNS activity, as indicated by SD1, compared with simply imagining one’s routine pre-performance scenario. As SD1 reflects the rest-and-digest functions of the PNS (Guyton & Hall, 2006; Saper, 2002), its lower values in the negative condition may indicate that participants were less relaxed before performing. This reduced relaxation may have led to decreased valence during performance, ultimately contributing to a lower perceived achievement.

Another possible explanation is that the no-memory condition imposed a lower cognitive load than either memory recall condition. We also observed that SD1 was higher in the no-memory condition than in the positive condition (**Figure 3E**), suggesting that imagining a familiar pre-performance state allowed the participants to remain more relaxed. Additionally, we did not control individual differences in what the participants envisioned during the no-memory condition. These factors may have enabled participants to achieve a relatively more comfortable state, leading to enhanced valence and, ultimately, improved performance compared to the negative condition.

### 4.4 Relationship between psychological and physiological indicators

After combining data from all conditions, the path analysis revealed that valence had a significant positive effect on performance achievement, and the SD2/SD1 ratio exerted a significant positive influence on valence (**Figure 4A**). However, SD2/SD1 did not directly predict performance achievement. These results suggest that while activation of the sympathetic nervous system does not directly enhance performance achievement, it does increase emotional valence, which, in turn, improves musical performance. Although the standardized estimate of SD2/SD1’s effect on valence was small, these findings support our hypothesis that sympathetic activity, when accompanied by positive emotions, contributes to enhanced performance achievement.

Conversely, SD1 did not significantly affect arousal or valence and negatively predicted performance achievement (**Figure 4A**). This indicates that PNS activity does not influence emotional arousal or valence and that an increase in SD1 may lead to lower performance achievement. As previously discussed, the PNS is associated with rest-and-digest functions (Guyton & Hall, 2006; Saper, 2002), which might seem beneficial for relaxation before performance. However, since moderate sympathetic activation is necessary to prepare the body for performance (Robazza et al., 2004; Yoshie et al., 2008b), these results underscore that relaxation through increased parasympathetic activity alone is insufficient for optimal performance. Instead, a balanced interaction between the sympathetic and parasympathetic nervous systems may be essential for achieving a high-level musical performance.

### 4.5 Correlation of subjective and objective evaluation

We found no significant differences in objective evaluations across the conditions (**Figure 5A**). However, there was a positive correlation between subjective performance achievement and objective evaluation (**Figure 5B**). Notably, the objective evaluations were based only on the first 20 seconds of the piece, with approximately six or seven notes. Despite this limited amount of information, the positive correlation suggests that the enhanced quality of performances, potentially arising from recalling autobiographical performance memories, was perceptible to the listeners. Future research should examine the criteria used in objective evaluations to determine which aspects of recorded performances are most effectively communicated. Understanding these criteria may offer insight into how positive autobiographical recall translates into perceptible improvements in musical output.

### 4.6 Limitation and future directions

This study had several limitations. First, because the experiment was conducted in a music studio, it remains unclear how these findings can be generalized to actual concert settings where professional musicians must avoid making mistakes. However, our supplementary analyses indicate that the quality of performance was effectively conveyed to listeners (Figure 5B), suggesting that the findings may hold true in real concert halls as well. To validate this possibility, future research should be conducted in concert halls with live audiences to provide and sophisticate a strategy to regulate sympathetic activation to enhance performance and realize full potential. This strategy may not be limited to professional musicians; it provides a new perspective for anyone required to present or express themselves in a public setting.

Second, ANS activity was assessed using HRV alone. Because HRV analysis requires several minutes of data, it is challenging to capture transient dynamic changes in ANS activity (Malik, 1996; Shaffer & Ginsberg, 2017). To achieve a more comprehensive and temporally sensitive evaluation, future studies should incorporate additional physiological indicators of ANS activity such as pupil diameter (Bishop et al., 2021) and skin conductance (Brouwer & Hogervorst, 2014). Including these measures may offer deeper insights into the dynamic ANS responses associated with recall of autobiographical performance memories and their influence on performance achievement.

## 5 Conclusion

We found that professional musicians who recalled memories of positive performance experienced greater performance achievement. Path analysis integrating data from all conditions revealed that, although SD2/SD1 did not directly predict performance achievement, increases in SD2/SD1 were associated with higher valence, which in turn led to improved achievement. These results suggest that consciously recalling positive memories before a performance in response to rising ANS activity may help professional musicians interpret their physiological state more positively, ultimately enhancing their on-stage performance.

## Data Availability Statement

The raw data supporting the conclusions of this article are available from the corresponding authors, without undue reservation.

## Author Contributions

Conceptualization: AW, TS, SF

Data Curation: AW, SK

Formal Analysis: AW, SK

Funding Acquisition: AW, SF

Investigation: AW

Methodology: AW, SK, TS, SF

Project Administration: SF

Software: AW

Supervision: SK, SF

Validation: AW, SK, TS

Visualization: AW, SF

Writing – Original Draft Preparation: AW

Writing – Review & Editing: AW, SK, TS, SF

## Acknowledgments

We are deeply grateful to Dr. Junichi Ushiyama for insightful comments on data analysis, and Dr. Michiko Yoshie for helpful comments on the experimental design. We also extend our heartfelt thanks to Mr. Motohiko Murabayashi, representative of the General Incorporated Foundation Runde, and Ms. Junko Shimamura for their generous support in providing access to the music studio.

